# Comprehensive interrogation of the ADAR2 deaminase domain for engineering enhanced RNA base-editing activity, functionality and specificity

**DOI:** 10.1101/2020.09.08.288233

**Authors:** Dhruva Katrekar, Nathan Palmer, Yichen Xiang, Anushka Saha, Dario Meluzzi, Prashant Mali

## Abstract

Adenosine deaminases acting on RNA (ADARs) can be repurposed to enable programmable RNA editing, however their exogenous delivery leads to transcriptome-wide off-targeting, and additionally, enzymatic activity on certain RNA motifs, especially those flanked by a 5’ guanosine is very low thus limiting their utility as a transcriptome engineering toolset. To address this, we explored comprehensive ADAR2 protein engineering via three approaches: First, we performed a novel deep mutational scan of the deaminase domain that enabled direct coupling of variants to corresponding RNA editing activity. Experimentally measuring the impact of every amino acid substitution across 261 residues, i.e. ~5000 variants, on RNA editing, revealed intrinsic domain properties, and also several mutations that greatly enhanced RNA editing. Second, we performed a domain-wide mutagenesis screen to identify variants that increased activity at *5’-GA-3*’ motifs, and discovered novel mutants that enabled robust RNA editing. Third, we engineered the domain at the fragment level to create split deaminases. Notably, compared to full-length deaminase overexpression, split-deaminases resulted in >1000 fold more specific RNA editing. Taken together, we anticipate this comprehensive deaminase engineering will enable broader utility of the ADAR toolset for RNA biotechnology and therapeutic applications.

## INTRODUCTION

Adenosine to inosine (A-to-I) editing is a common post-transcriptional modification in RNA that occurs in a variety of organisms, including humans. This A-to-I deamination of specific adenosines in double-stranded RNA is catalyzed by enzymes called adenosine deaminases acting on RNA (ADARs) (*1–12*). Since inosine is structurally similar to guanosine, it is interpreted as a guanosine during the cellular processes of translation and splicing, thereby making ADARs powerful systems for altering protein sequences.

Correspondingly, adenosine deaminases have been repurposed for site-specific RNA editing by recruiting them to target RNA sequences using engineered ADAR-recruiting RNAs (adRNAs) (*13*). Recently, several studies have demonstrated the potential of both genetically encodable and chemically modified RNA-guided adenosine deaminases for the correction of point mutations and the repair of premature stop codons both *in vitro* (*14–22*) and *in vivo 23, 24*. These studies have primarily relied on exogenous ADARs which introduce a significant number of transcriptome wide off-target A-to-I edits (*16, 23, 25, 26*). One solution to this problem is the engineering of adRNAs to enable the recruitment of endogenous ADARs. In this regard, we recently showed that using simple long antisense RNA (>60bp) can suffice to recruit endogenous ADARs and these adRNAs are both genetically encodable and chemically synthesizable *(23)*; and Merkle and colleagues showed that using engineered chemically synthesized antisense oligonucleotides (*18*) could also lead to robust RNA editing via endogenous ADAR recruitment. Although this modality allows for highly specific editing, its applicability is restricted to editing adenosines in certain RNA motifs preferred by the native ADARs, and in tissues with high endogenous ADAR activity. Additionally, it cannot be utilized for novel functionalities such as deamination of cytosine to uracil (C-to-U) editing which requires exogenous delivery of ADAR2 variants (*27*). Thus, engineering a genetically encodable RNA-editing tool that efficiently edits RNA with high specificity and activity is essential for enabling broader use of this toolset for biotechnology and therapeutic applications.

In this regard, the crystal structure of the ADAR2 deaminase domain (ADAR2-DD) (*28–30*) and several pioneering biochemical and computational studies (*31–40*) have laid the foundation for understanding its catalytic mechanism and target preferences, but we still lack comprehensive knowledge of how mutations and fragmentation affect the ability of the ADAR2-DD to edit RNA. To address this, we first carried out a quantitative deep mutational scan (DMS) of the ADAR2-DD, measuring the effect of every possible point mutation on enzyme function. We utilized the sequence-function map thus generated, to identify novel enhanced variants for A-to-I editing. Additionally, combining information from these sequence-function maps with existing knowledge of the structure and residue conservation scores, we also engineered a genetically encodable split-ADAR2 system that enabled efficient and highly specific RNA editing.

## RESULTS

### Deep mutational scanning of the ADAR2 deaminase domain

To gain comprehensive insight into how mutations affect the ADAR2 deaminase domain (ADAR2-DD), we used deep mutational scanning (DMS), a technique that enables simultaneous assessment of the activities of thousands of protein variants (*41, 42*). Typically, this approach relies on phenotypic selection methods such as cell fitness or fluorescent reporters that result in an enrichment of beneficial variants and a depletion of deleterious variants. However, as RNA editing yields are not precisely quantifiable using surrogate readouts, we focused on directly measuring enzymatic activity in the screens. To do so, we linked genotype to phenotype by placing the RNA editing site on the same transcript encoding the deaminase variant, and ensuring every cell in the pooled screen received a single library element. This novel approach enabled us to perform a quantitative deep mutational scan of the core 261 amino acids (residues 340-600) of the ADAR2-deaminase domain via 4959 (261×19) single amino acid variants, measuring the effect of each mutation on adenosine to inosine (A-to-I) editing yields (**Figure 1a**).

**Figure 1.**
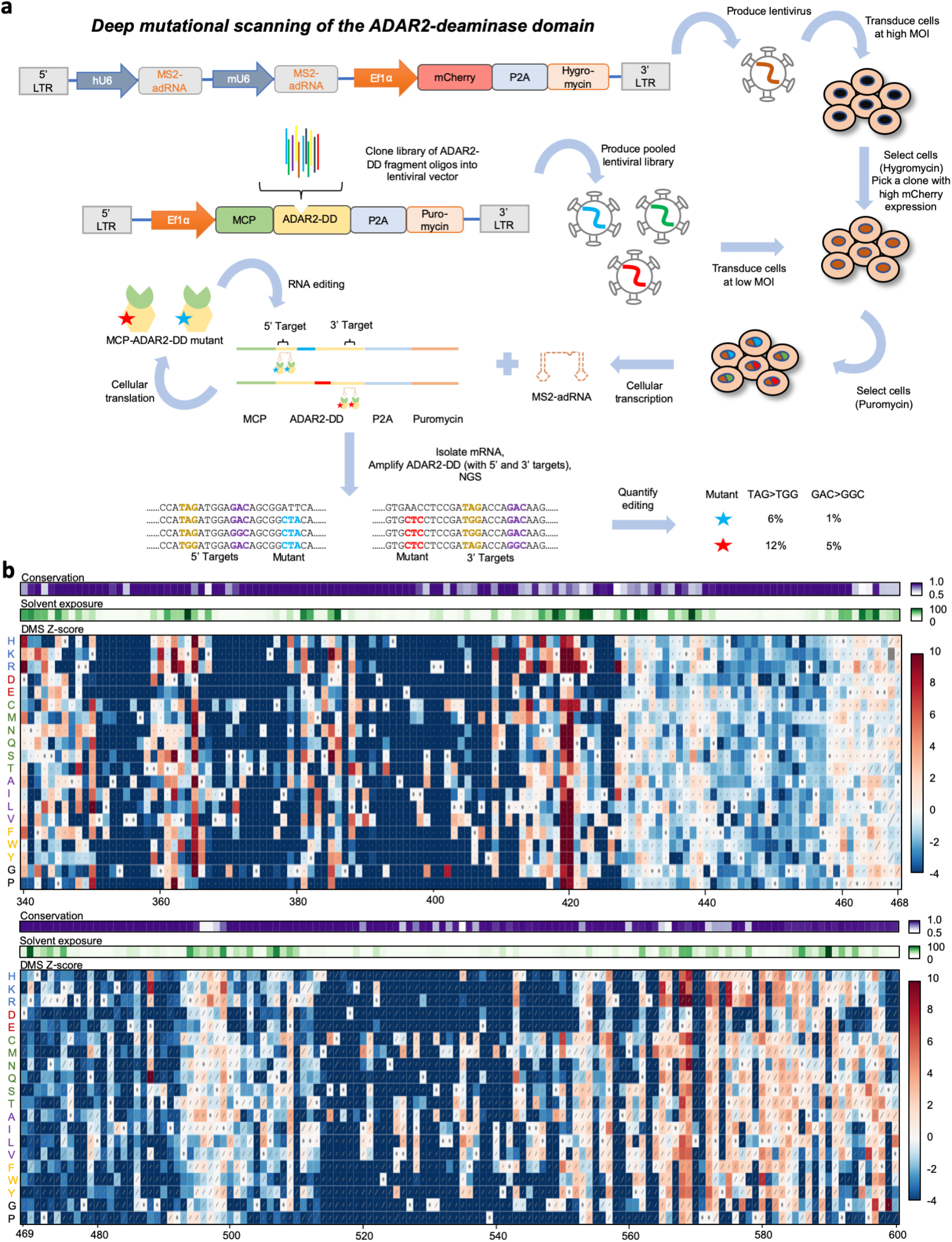
(**a**) Schematic of the deep mutational scanning approach. HEK293FT cells were transduced with the MS2-adRNA lentiviruses at a high MOI and a single clone was selected based on mCherry expression. These cells bearing the MS2-adRNA were then transduced with the lentiviral library of MCP-ADAR2-DD-NES variants at a low MOI to ensure delivery of a single variant per cell. Upon translation in the cell, each MCP-ADAR2-DD variant, in combination with the MS2-adRNA, edited its own transcript creating a synonymous change. These transcripts were then sequenced to quantify the editing efficiency associated with each variant. (**b**) Heatmaps illustrating impact of single amino acid substitutions in residues 340-600 on the ability of the ADAR2-DD to edit a UAG motif. Rectangles are colored according to the scale bar on the right depicting the Z-score for editing a UAG motif as compared to the ADAR2-DD. Diagonal bars indicate standard error. The amino acids in the wild-type ADAR2-DD are indicated in the heatmap with a ∙. Amino acids are indicated on the left and grouped based on type of amino acid: positively charged, negatively charged, polar-neutral, non-polar, aromatic and unique. The heatmap bars at the top represent amino acid conservation score and surface exposure respectively.

Given the large size of the deaminase domain at >750bp, the library was created using 6 tiling oligonucleotide pools (**Supplementary Figure 1a**). These pools were cloned into a lentiviral vector containing the MS2 coat protein (MCP) and the remainder of the deaminase domain and a puromycin resistance gene (**Figure 1a, Supplementary Figure 1b**). Editing sites were chosen within the deaminase domain, outside of the mutated residues, such that an A-to-I change would result in a synonymous mutation. To ensure read length coverage in next generation sequencing, members of the first three library pools were assayed for editing at the 5’ end while the remaining members were assayed at the 3’ end of the deaminase domain (**Supplementary Figure 1a**). Towards this, two HEK293FT clonal cell lines were created with MS2-adRNAs targeting 5’ and 3’ UAG sites integrated into them. The scan was carried out in cell lines harboring these MS2-adRNAs by transducing them with the corresponding libraries at a low MOI (0.2-0.4). Following lentiviral transduction and puromycin selection, RNA was extracted from the harvested cells and reverse transcribed. Relevant regions of the deaminase domain were amplified from the cDNA and sequenced (**Supplementary Figure 1c**). 4958 of the 4959 possible variants were successfully detected. The deaminase domain transcripts for each variant also contained the associated A-to-I editing yields, which were then quantified for both replicates of the DMS (**Supplementary Figure 1d**).

The scans revealed both intrinsic domain properties, and also several mutations that enhanced RNA editing (**Figures 1b, 2a, Supplementary Figure 2a**). Specifically: 1) As expected, most mutations in conserved regions 442-460 and 469-495 that bind the RNA duplex near the editing site led to a significant decrease in editing efficiency of the enzyme (*29*); 2) However, mutating the negatively charged E488 residue, which recognizes the cytosine opposite the flipped adenosine by donating hydrogen bonds, to a positively charged or most polar-neutral amino acids resulted in an improvement in editing efficiency. This is consistent with the previously discovered E488Q mutation which has been shown to improve the catalytic activity of the enzyme (*32*); 3) Furthermore, most mutations to residues that contact the flipped adenosine (V351, T375, K376, E396, C451, R455) were observed to be detrimental to enzyme function (*29*); 4) Similarly, the residues of the ADAR2-DD that interact with the zinc ion in the active site and the inositol hexakisphosphate (R400, R401, K519, R522, S531, W523, D392, K483, C451, C516, H394 and E396) were all also extremely intolerant to mutations (*28*). 5) Additionally, as expected, surface exposed residues in general readily tolerated mutations as compared to buried residues (*29*).

**Figure 2.**
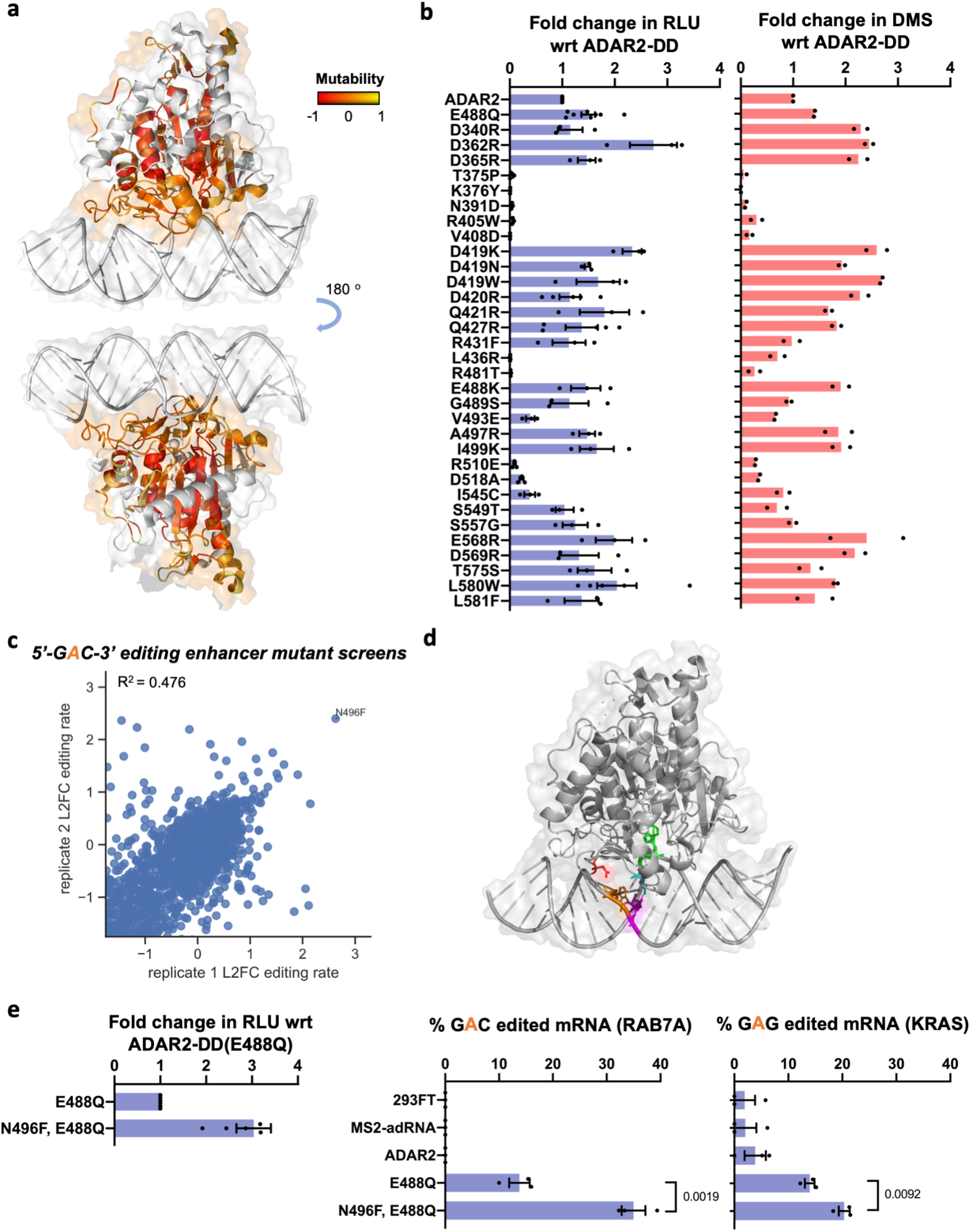
(**a**) Structure of the ADAR2-DD bound to its substrate (PDB 5HP3) with the degree of mutability of each residue as measured by the DMS highlighted. Residues that are highly intolerant to mutations are colored red while residues that are highly mutable are colored yellow. Residues not assayed in this DMS are colored white. (**b**) List of mutants from the pooled DMS screens were individually validated in an arrayed luciferase assay using a cluc reporter bearing a UAG stop codon. The plots represent fold change as compared to the wild-type ADAR2 for (i) the arrayed luciferase assay and (ii) the DMS screen. Values represent mean +/− SEM for the luciferase assay (n>2) and mean for the DMS (n=2). (**c**) Using the library chassis of the DMS, a screen of deaminase domain mutants (in an E488Q background) was performed to mine variants with improved activity against *5’-GA-3’* RNA motifs. (**d**) Structure of the ADAR2-DD(E488Q) bound to its substrate (PDB 5ED1) with the N496 residue highlighted in red, the E488Q residue in cyan, the target adenosine in green, the orphaned cytosine in magenta and the adenosine on the unedited strand that base pairs with the 5’ uracil flanking the target adenosine in orange. (**e**) (i) The N496F, E488Q mutant was validated in a luciferase assay using a cluc reporter bearing a UGA stop codon. The plot represents fold change as compared to the ADAR2-DD(E488Q). Values represent mean +/− SEM (n=6). (ii) Editing of a GAC motif in the 3’UTR of the RAB7A transcript, and (iii) a GAG motif in the CDS of the KRAS transcript. Values represent mean +/− SEM (n=3). All experiments were carried out in HEK293FT cells.

To independently validate the results from the DMS, we individually examined 33 mutants from the DMS whose editing efficiencies ranged from very low to very high as compared to the wild-type ADAR2-DD. The mutants were assayed for their ability to repair a premature amber stop codon (UAG) in the *cypridina* luciferase (cluc) transcript (*16*). We observed that a majority of the mutants (85%) followed the same trend in our arrayed validations as seen in the pooled screens (**Figure 2b**). Additionally, we compared the efficiency of variants in our ADAR2-DD DMS at editing UAG triplets, to published mutants (*29, 30, 32*) and again observed similar agreement in the activity of a majority of the variants (75%), together confirming the efficacy of the deep mutational scan.

### Enhancing functionality of the ADAR2 deaminase domain

Building on this platform (**Figure 1a**), we next screened for domain variants that expanded its functionality, in particular focusing on mining mutants that improved editing at refractory RNA motifs such as adenosines flanked by a 5’ guanosine (*26, 32*). Towards this, two HEK293FT clonal cell lines were created with MS2-adRNAs targeting 5’ and 3’ GAC sites integrated into them. A screen was carried out in cell lines harboring these MS2-adRNAs by transducing them with the corresponding MCP-ADAR2-DD(E488Q) libraries at a low MOI (0.2-0.4), evaluating the potential of 3287 mutants to edit a GAC motif. Similar to above, following lentiviral transduction and selection, RNA was extracted, reverse transcribed, and relevant regions of the deaminase domain amplified, sequenced and analyzed (**Figure 2c**). Via this, we discovered a novel mutant N496F that enhanced editing at a *5’-GA-3’* motif. Interestingly, in the ADAR2-DD crystal structure, the N496 residue is in close proximity to the adenosine on the unedited strand that base pairs with the 5’ uracil flanking the target adenosine (**Figure 2d**) (*29*). We validated this mutant using a cluc luciferase reporter bearing a premature opal stop codon (UGA) and confirmed that the N496F, E488Q double mutant was 3-fold better at restoring luciferase activity as compared to E488Q alone (**Figure 2e**). To further confirm that the N496F, E488Q double mutant could be used to efficiently edit adenosines flanked by a 5’ guanosine, we tested the ability of this mutant to edit a GAC and GAG motif in the 3’ UTR and CDS of the endogenous RAB7A and KRAS transcripts respectively. We observed that the double mutant N496F, E488Q was 2.5-fold more efficient at editing the GAC motif and 1.5-fold more efficient at editing a GAG motif than the E488Q (**Figure 2e, Supplementary Figure 3**), together confirming the ability of this novel screening format to discover variants that expand the deaminase domain functionality.

### Improving specificity via splitting of the ADAR2 deaminase domain

In addition to increasing the on-target activity of ADARs at editing adenosines in non-preferred motifs, another challenge towards unlocking their utility as a RNA editing toolset is that of improving specificity. Due to their intrinsic dsRNA binding activity, overexpression of ADARs leads to promiscuous transcriptome wide off-targeting, and thus, when relying on exogenous ADARs, it is important to engineer restriction of the catalytic activity of the overexpressed enzyme only to the target mRNA. We hypothesized that it might be possible to achieve this by splitting the deaminase domain into two catalytically inactive fragments that come together to form a catalytically active enzyme only at the intended target (**Figure 3a**). Since we and others have utilized the MS2 Coat Protein (MCP) and Lambda N (λN) systems to efficiently recruit ADARs, we first decided to utilize these systems to recruit the two split halves, i.e. the N- and C-terminal fragments of the ADAR2-DD (*14, 23*). Specifically, constructs were created with cloning sites for N-terminal fragments located downstream of the MCP while those for the C-terminal fragments located upstream of the λN. Chimeric adRNAs were designed to bear a BoxB and a MS2 stem loop along with an antisense domain complementary to the target. Studying the sequence-function map of the ADAR2-DD generated from the DMS (**Figure 1b**) as well as its crystal structure we identified 18 putative regions for splitting the protein (**Figure 3b**) (*29, 43*). The resulting 18 different split-ADAR2 pairs were assayed for their ability to repair a premature amber stop codon (UAG) in the *cypridina* luciferase (cluc) transcript in the presence of the recruiting adRNA bearing BoxB and MS2 stem loops (**Figure 3c**). Of these pairs 9-12 showed the best editing efficiency, and notably were all located within residues 465-468 which have low conservation scores across species (*29*). Interestingly, this region is flanked by highly conserved amino acids (442-460 and 469-495).

**Figure 3.**
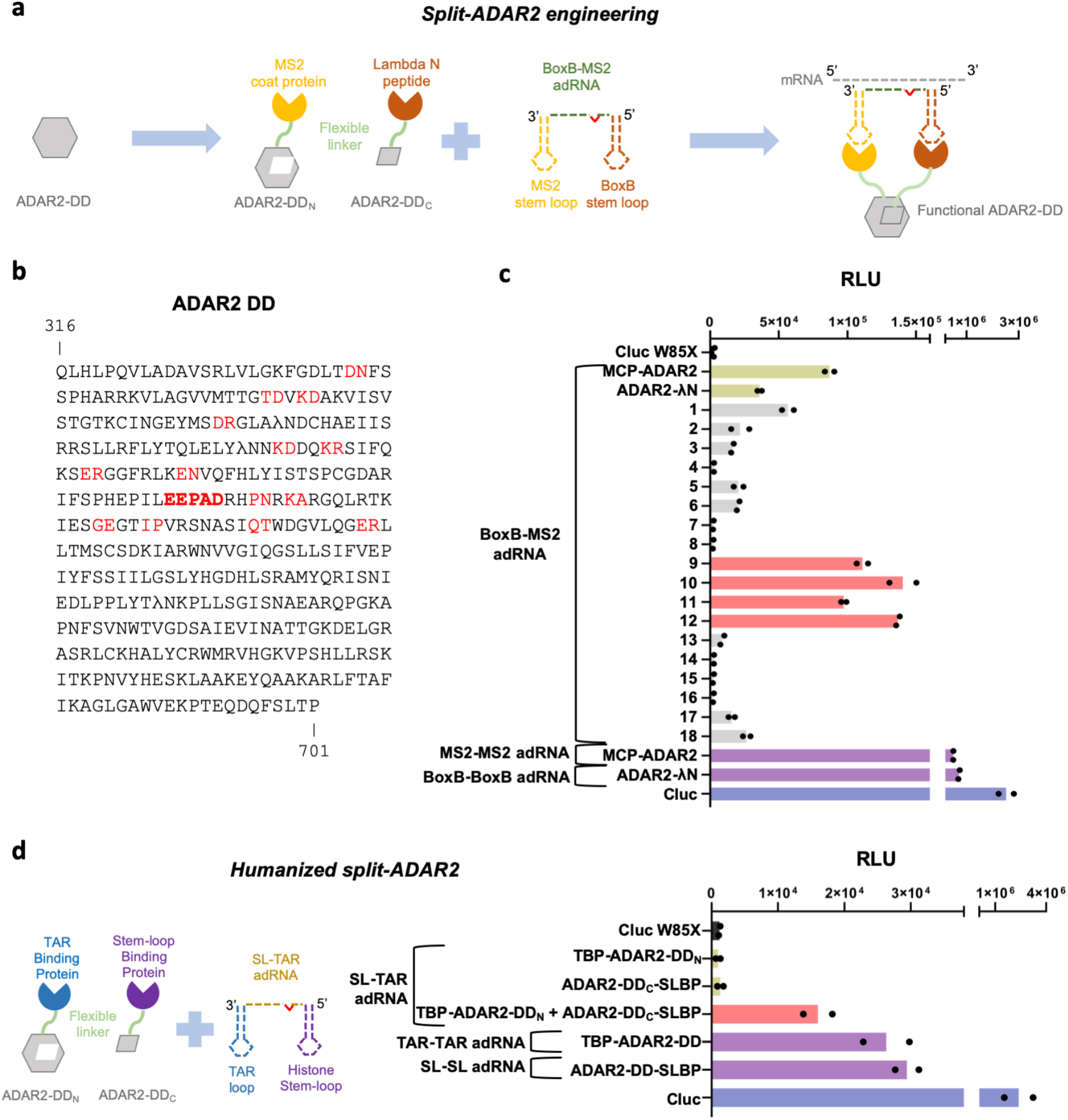
(**a**) Schematic of the split-ADAR2 engineering approach. (**b**) Sequence of the ADAR2-DD. The protein was split between residues labelled in red, and a total of 18 pairs were evaluated. (**c**) The ability of each split pair from (b) to correct a premature stop codon when transfected with a chimeric BoxB-MS2 adRNA was assayed via a luciferase assay. The pairs 1-18 correspond to the residues in red in (b) in the order in which they appear. The residues in (b) in bold red correspond to pairs 9-12. Values represent mean (n=2). (**d**) Engineering of humanized split-ADAR2 variant based on pair 12 and assayed of its ability to correct a stop codon in the cluc transcript. Values represent mean (n=2). All experiments were carried out in HEK293FT cells.

We also confirmed that every component of the split-ADAR2 system was essential for RNA editing. Specifically, we assayed all components and pairs of components for their ability to restore luciferase activity. The MCP-ADAR2-DD was included as a control. We observed restoration of luciferase activity only when every component of the split-ADAR2 system was delivered, confirming that the individual components lacked enzymatic activity (**Supplementary Figure 4a**). Additionally, we also confirmed the importance of fragment orientation for the formation of a functional enzyme. Towards this, we swapped the positions of the N- and C-terminal fragments and created ADAR2-DD_N_-MCP and λN-ADAR2-DD_C_ in addition to the working MCP-ADAR2-DD_N_ and ADAR2-DD_C_-λN pair. We then tested each pair of N- and C-terminal fragments and observed functionality only for the MCP-ADAR2-DD_N_ paired with ADAR2-DD_C_-λN (**Supplementary Figure 4b**).

Since MCP and λN are proteins of viral origin we next replaced these with the human TAR Binding Protein (TBP) and the Stem Loop Binding Protein (SLBP) respectively to create a humanized split-ADAR2 system with improved translational relevance (*44*). In the presence of a chimeric adRNA containing a histone stem loop and a TAR stem loop, we observed restoration of luciferase activity (**Figure 3d**). This also confirmed that the split-ADAR2 pair 12 (hereon referred to as ADAR2-DD_N_ and ADAR2-DD_C_) could indeed be recruited for RNA editing using two independent sets of protein-RNA binding systems.

Finally, we investigated the specificity profiles via analysis of the transcriptome-wide off-target A-to-G editing effected by this system (**Figure 4a, 4b** and **Supplementary Figures 5, 6**). Each condition from **Figure 4a** (where the endogenous RAB7A transcript was targeted) was analyzed by RNA-seq. From each sample, we collected ~19 million uniquely aligned sequencing read pairs. We then used Fisher’s exact test to quantify significant changes in A-to-G editing yields, relative to untransfected cells, at each reference adenosine site having sufficient read coverage. Notably, utilizing the split-ADAR2 system observed a 1100-1400 fold reduction in the number of off-targets as compared to the MCP-ADAR2 system. Excitingly, the specificity profiles of the split-ADAR2 system were comparable to those seen when using endogenous recruitment of ADARs via long antisense RNA (*23*) (**Supplementary Figures 5, 6**).

**Figure 4.**
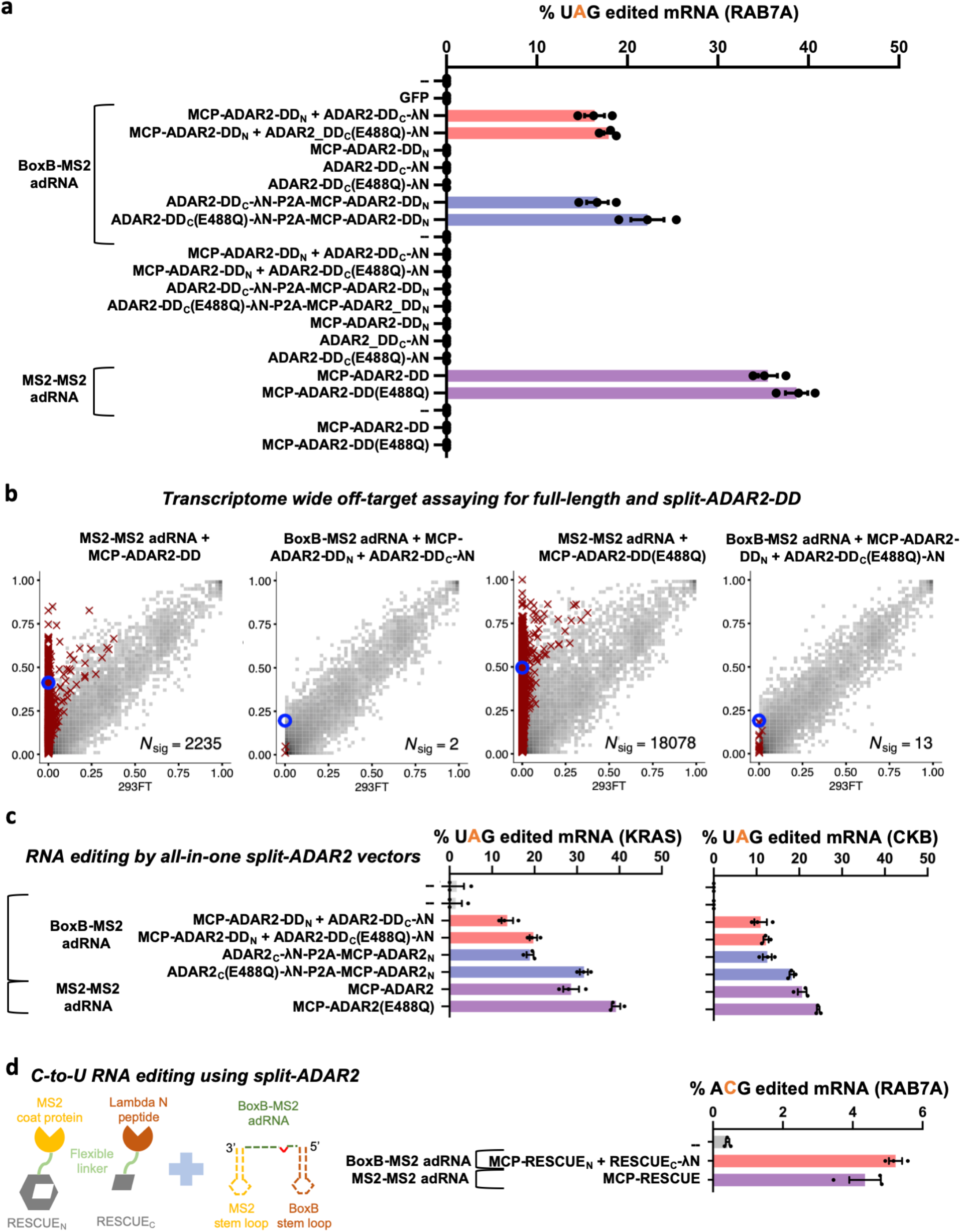
(**a**) The components of the split-ADAR2 system based on pair 12 were tested for their ability to edit the RAB7A transcript. Editing was observed only when every component was delivered. Values represent mean +/− SEM (n=3). (**b**) 2D histograms comparing the transcriptome-wide A-to-G editing yields observed with each construct (*y*-axis) to the yields observed with the control sample (*x*-axis). Each histogram represents the same set of reference sites, where read coverage was at least 10 and at least one putative editing event was detected in at least one sample. Bins highlighted in red contain sites with significant changes in A-to-G editing yields when comparing treatment to control sample. Red crosses in each plot indicate the 100 sites with the smallest adjusted *P* values. Blue circles indicate the intended target A site within the RAB7A transcript. All experiments were carried out in HEK293FT cells. (**c**) The split-ADAR2 system was assayed for editing the KRAS and CKB transcripts. Values represent mean +/− SEM (n=3). (**d**) A split-RESCUE was engineered based on pair 12 and assayed for C-to-U editing of the RAB7A transcript. Values represent mean +/− SEM (n=3).

To confirm generalizability of the results, we also tested the split-ADAR2 at two additional endogenous loci: an adenosine in the 3’UTR of CKB and an adenosine in the CDS of KRAS, and observed robust editing efficiency of the split-ADAR2 system (**Figure 4a, 4c**). To enable convenient delivery of the split-ADAR2 system we also created an all-in-one vector bearing a bicistronic ADAR2-DD_C_-λN-P2A-MCP-ADAR2-DD_N_ which also enabled higher editing efficiencies across all three loci tested (**Figures 4a, 4c**). The entire split-ADAR2 system consisting of CMV promoter driven ADAR2-DD_C_-λN-P2A-MCP-ADAR2-DD_N_ and a human U6 promoter driven BoxB-MS2 adRNA is ~3500 bp in size and can easily be packaged into a single adeno-associated virus (AAV).

Lastly, to test if the split-ADAR2 chassis could be expanded to enable new functionalities, specifically C-to-U editing, we used it create a split-RESCUE system and confirmed comparable C-to-U RNA editing of the endogenous RAB7A transcript as the full-length MCP-RESCUE (*27*) (**Figure 4d**).

## DISCUSSION

Towards addressing two of the fundamental challenges in using ADARs for programmable RNA editing, specifically, one, exogenous delivery leading to massive transcriptome-wide off-targeting (*16, 23, 25, 26*), and two, poor enzymatic activity on certain RNA motifs such as those flanked by a 5’ guanosine (*26, 32*), we have explored in this study comprehensive ADAR2 deaminase protein engineering via three distinct approaches. First, we performed a novel deep mutational scan, comprehensively assaying all possible single amino acid substitutions of 261 residues of the deaminase domain for their impact on RNA editing yields. We created a sequence-function map of the deaminase domain that complements existing knowledge derived from prior structure and biochemistry-based studies and improves our understanding of the enzyme, and can serve as a map for engineering novel variants with tailored activity for specific applications. Second, we used this novel screening chassis to also expand deaminase functionality by performing a domain-wide mutagenesis screen to identify variants that increased activity at *5’-GA-3*’ motifs, and discovered novel variants that enabled robust RNA editing such as ADAR2-DD(N496F, E488Q). Specifically, this mutant was 1.5-2.5 fold more efficient at editing adenosines with a 5’ guanosine than the classic hyperactive ADAR2-DD(E488Q). Finally, third, we engineered the deaminase domain at the fragment level to create split deaminases each of which was inactive by itself but together formed a functional enzyme upon combining at the target site. This split-ADAR2 tool was highly transcript specific (>1000 fold compared to full domain over expression), and with off-target profiles similar to those seen via recruitment of endogenous ADARs (*23*). We believe that creation of the split-ADAR2 tool paves the way for the use of the highly active ADAR2 deaminase domain variants discovered in our deep mutational scans towards enabling broader utility of the ADAR toolset for biotechnology and therapeutic applications. Additionally, these approaches could also be applied to the study and engineering of other RNA modifying enzymes (*45, 46*).

## Acknowledgements

We thank members of the Mali lab for discussions, advice and help with experiments. This work was generously supported by UCSD Institutional Funds and NIH grants (R01HG009285, RO1CA222826, RO1GM123313, 1K01DK119687). This publication includes data generated at the UC San Diego IGM Genomics Center utilizing an Illumina NovaSeq 6000 that was purchased with funding from a National Institutes of Health SIG grant (#S10 OD026929).

## Author contributions

D.K. and P.M. conceived the study and wrote the paper. D.K., Y.X., P.M., and A.S. performed experiments. N.P. analyzed the deep mutational scan. D.M. quantified RNA-editing activity from RNA-seq data.

---

**SI Figure 1:**
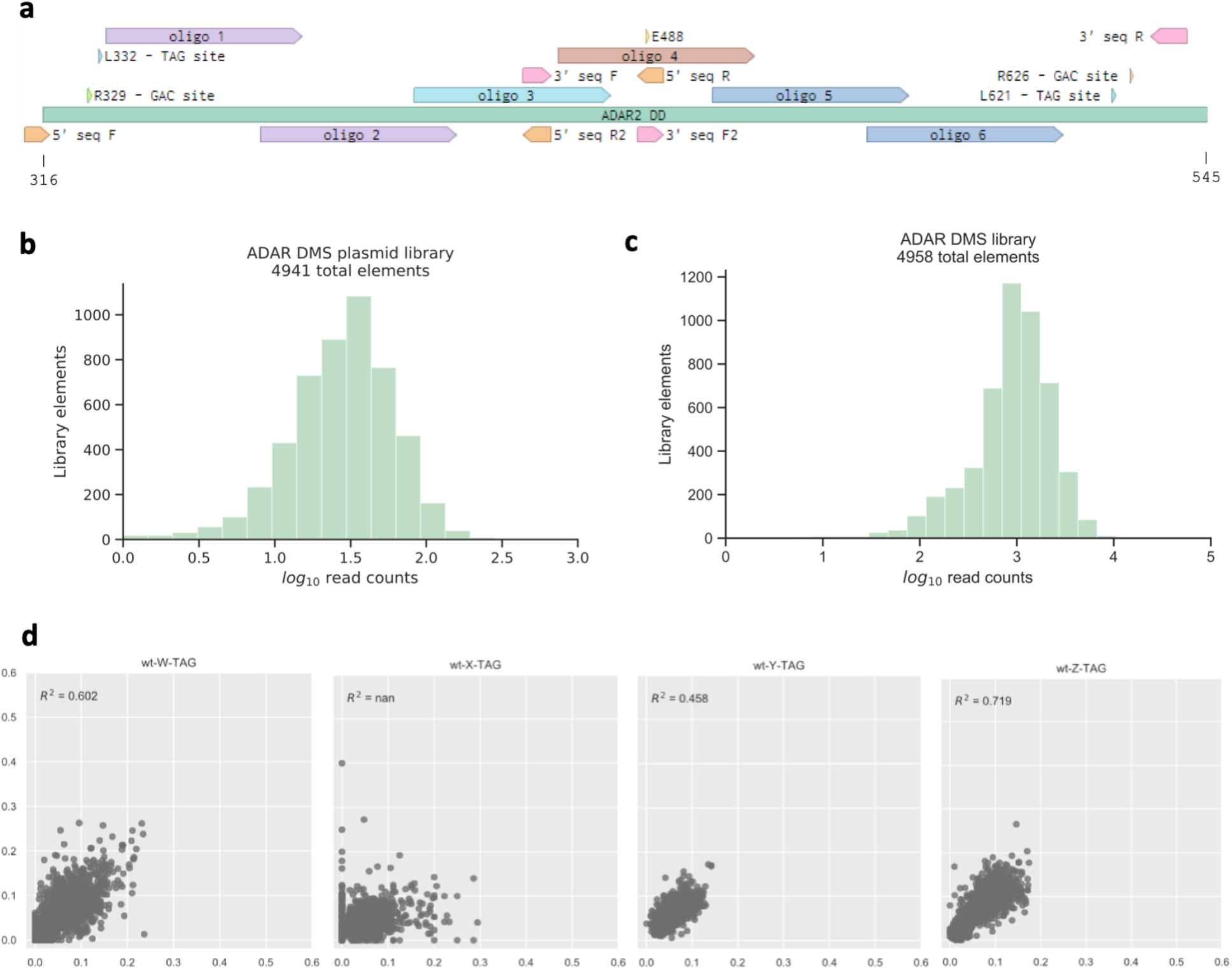
(**a**) Schematic of the ADAR2-DD showing oligonucleotide pools used to create the DMS library along with editing sites and primer binding sites. Oligonucleotide libraries 1, 2 and 3 were assayed for editing at the sites located at the 5’ end while libraries 4, 5 and 6 were assayed for editing at the 3’ end. Libraries 1 and 2 were amplified using primers 5’ seq F and 5’ seq R2, library 3 with 5’ seq F and 5’ seq R, library 4 with 3’ seq F and 3’ seq R and libraries 5 and 6 with 3’ seq F2 and 3’ seq R. (**b**) Library coverage of the ADAR2-DD DMS plasmids. (**c**) Histogram of variant counts from the DMS. 4958 of the 4959 variants were detected. (**d**) Replicate correlation for the ADAR2-DD DMS. The X and Y axes on every plot represent the fraction of edited reads.

**SI Figure 2:**
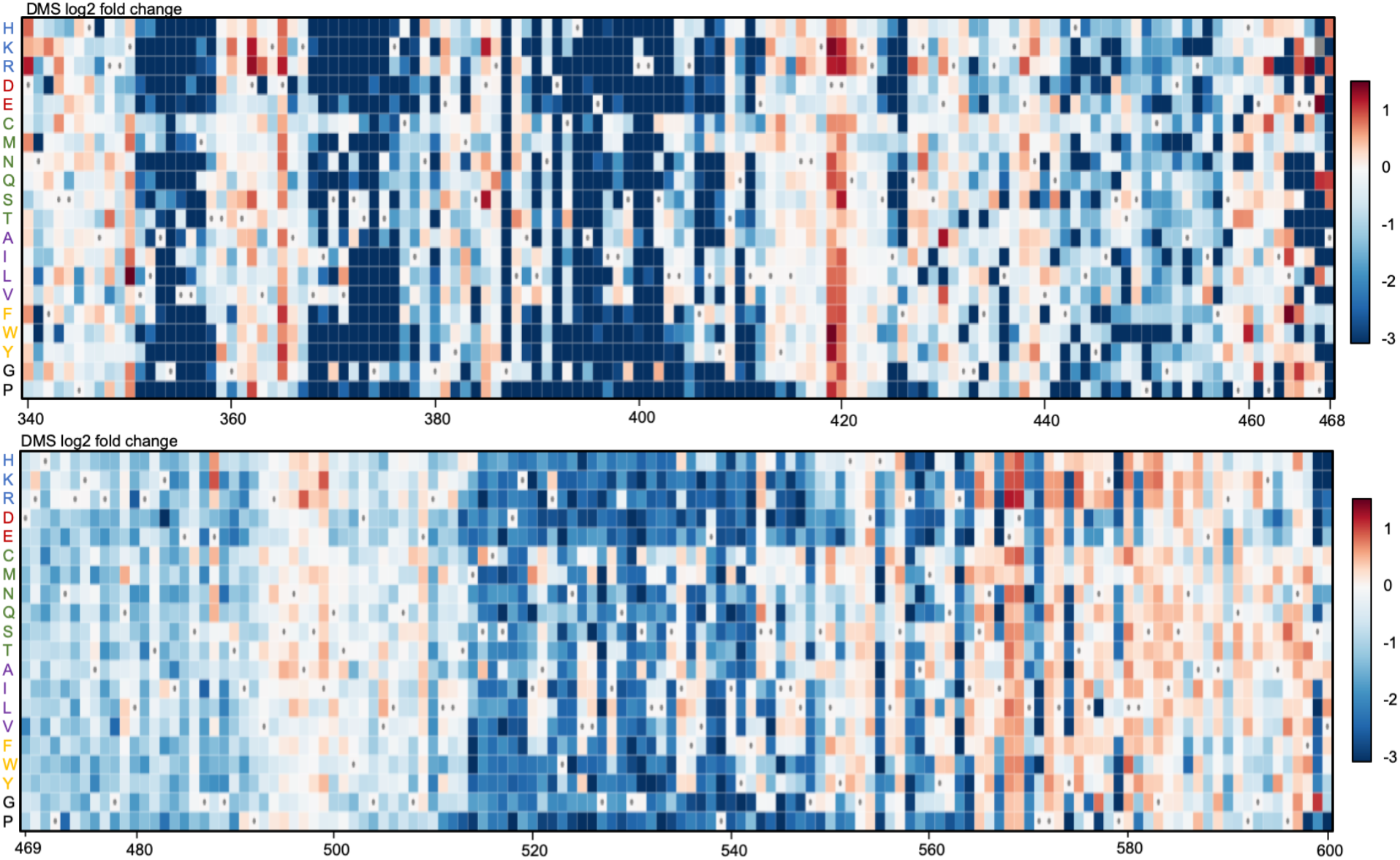
Heatmaps illustrating how single amino acid substitutions in residues 340-600 impact the ability of the ADAR2-DD to edit a UAG motif. Rectangles are colored according to the scale bar on the bottom right depicting the geometric mean of log2 fold change in editing efficiency as compared to the ADAR2-DD. The amino acids in the wild-type ADAR2-DD are indicated in the heatmap with a ∙. Amino acids are indicated on the left and grouped based on type of amino acid: positively charged, negatively charged, polar-neutral, non-polar, aromatic and unique.

**SI Figure 3:**
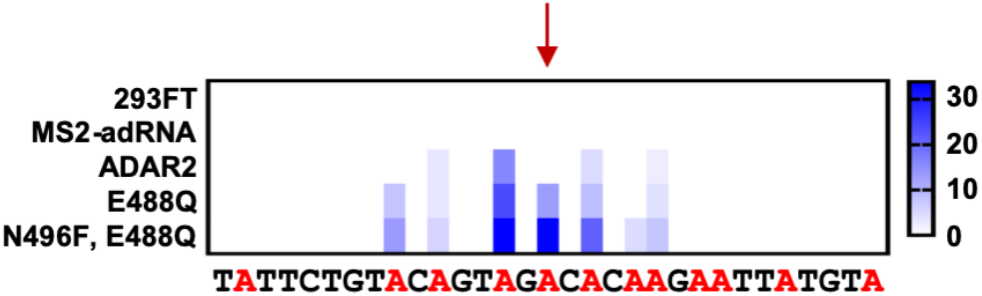
Heatmap depicting hyper-editing observed with the N496F, E488Q double mutant corresponding to the RAB7A plot in Figure 2e. The red arrow indicates the target.

**SI Figure 4:**
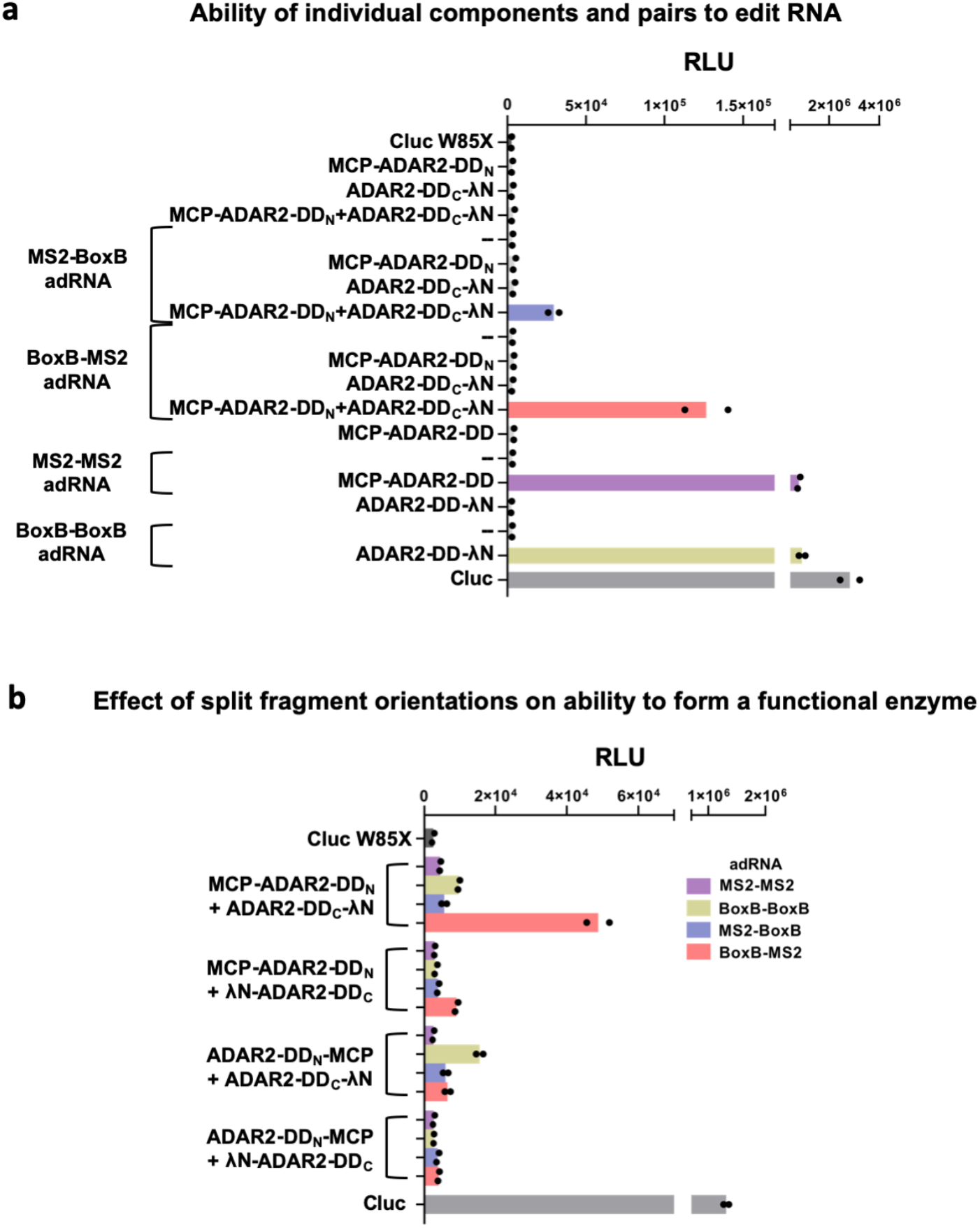
(**a**) All components of the split-ADAR2 system were tested for their ability to edit RNA via the luciferase assay. Restoration of luciferase activity is observed only when every component is delivered. Values represent mean (n=2). (**b**) The importance of orientation of the N- and C-terminal fragments in forming a functional ADAR2-DD is assayed via the luciferase assay. Chimeric and non-chimeric adRNA are used to recruit the split-ADAR2 pairs. Values represent mean (n=2).

**SI Figure 5:**
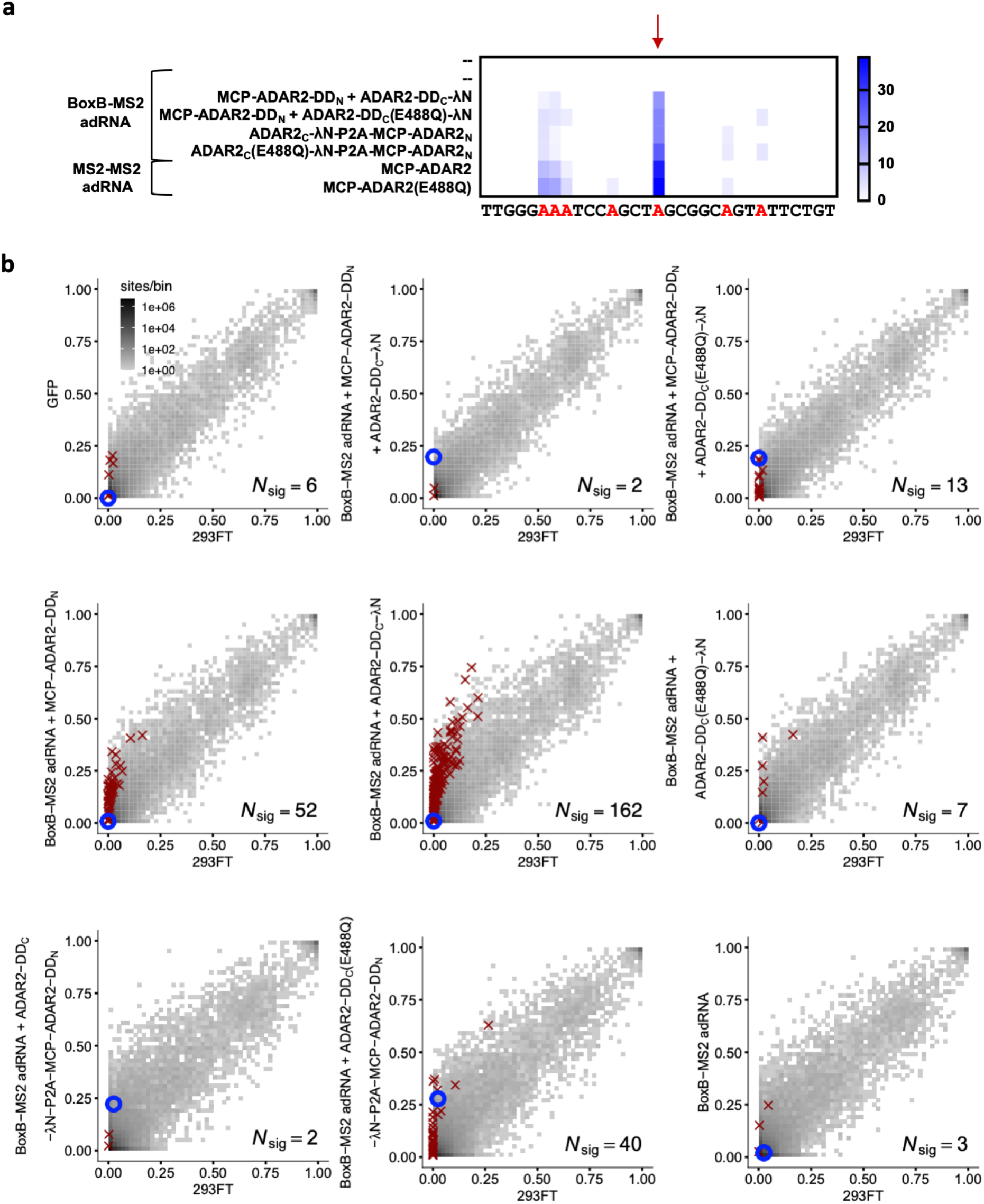
(**a**) Heatmap depicting hyper-editing observed with the split-ADAR2 system corresponding to the plot in Figure 4a. The red arrow indicates the target adenosine. (**b**) 2D histograms comparing the transcriptome-wide A-to-G editing yields observed with each construct from Figure 4a (y-axis) to the yields observed with the control sample (x-axis). Each histogram represents the same set of 22583 reference sites, where read coverage was at least 10 and at least one putative editing event was detected in at least one sample. Bins highlighted in red contain sites with significant changes in A-to-G editing yields when comparing treatment to control sample. Red crosses in each plot indicate the 100 sites with the smallest adjusted p-values. Blue circles indicate the intended target A-site within the RAB7A transcript. Large counts in bins near the lower-left corner likely correspond not only to low editing yields in both test and control samples, but also to sequencing errors and alignment errors. Large counts in bins near the upper-right corner of each plot likely correspond to homozygous single nucleotide polymorphisms (SNPs), as well as other differences between the reference genome and the genome of the HEK293FT cell line used in the experiments.

**SI Figure 6:**
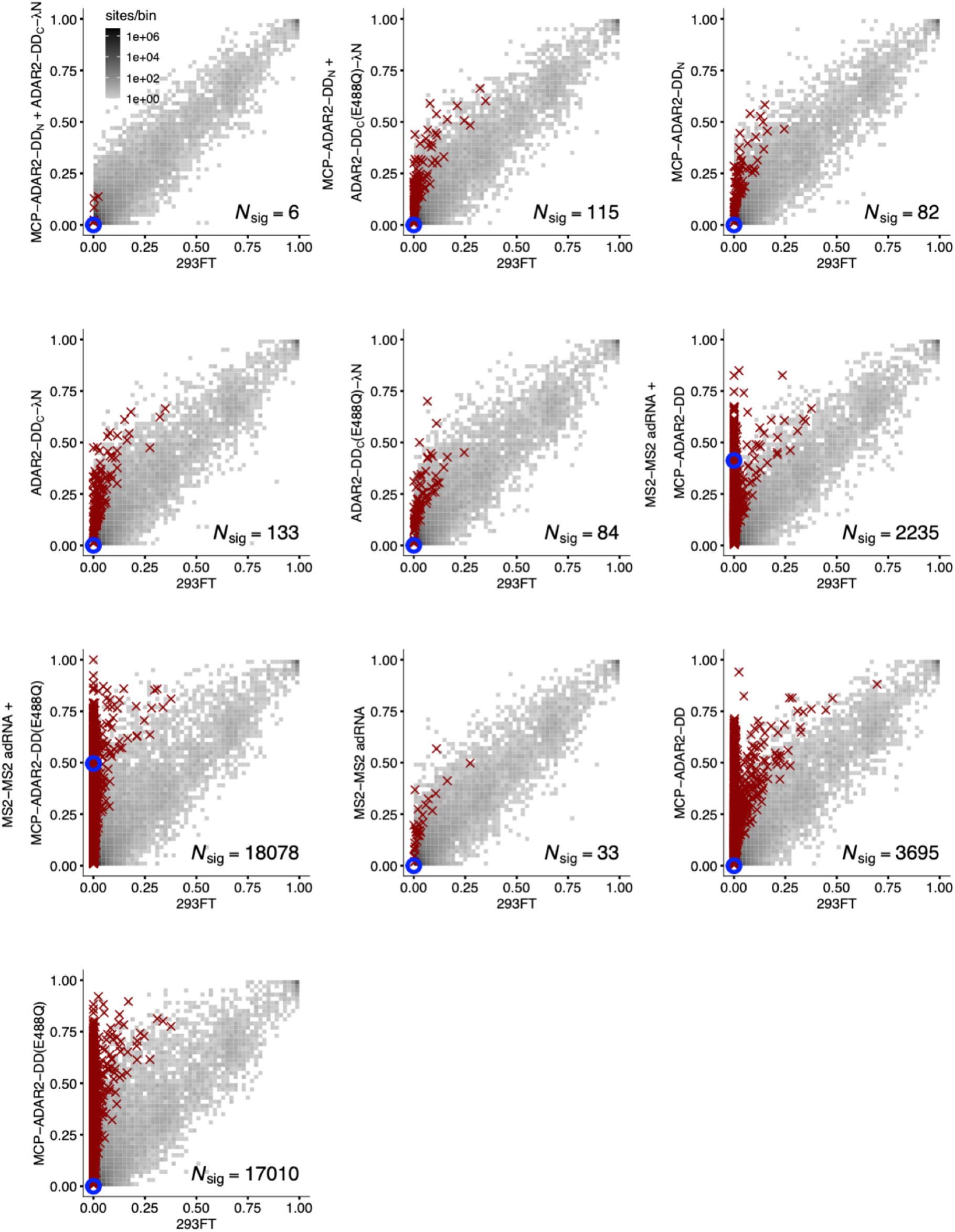
2D histograms comparing the transcriptome-wide A-to-G editing yields observed with each split-ADAR2 construct (y-axis) to the yields observed with the control sample (x-axis).

## METHODS

### Deep mutational scan and screen

#### i. Oligonucleotide pools

To create the library of single amino acid substitutions in the ADAR2 deaminase domain, we ordered an oligonucleotide chip (CustomArray) consisting of 6 oligonucleotide pools (each 168 bp in length). These pools, in combination, spanned residues 340-600 of the ADAR2 deaminase domain. Each of these pools was amplified in a 50 μl PCR reaction using Kapa HiFi HotStart PCR Mix (Kapa Biosystems), 40 ng of synthesized oligonucleotide as template and pool-specific primers. The 6 PCR products were purified using the QIAquick PCR Purification Kit (Qiagen) to eliminate byproducts.

#### ii. Creation of vectors for cloning oligonucleotide pools

We ordered a gene block (IDT) for MCP-ADAR2-DD-NES and used mutagenesis PCR to create the MCP-ADAR2-DD(E488Q)-NES. These fragments were then used as templates to generate 6 PCR fragments from which deletions of the MCP-ADAR2-DD-NES and the MCP-ADAR2-DD(E488Q)-NES were created. The deleted regions corresponded to the sequence covered by each of the 6 oligonucleotide pools and was replaced instead with an Esp3I digestion site. To create the plasmid library, we began by mutating the two Esp3I digestion sites in the LentiCRISPR v2 plasmid (gift from Feng Zhang, Addgene #52961)(*47*) using PCR mutagenesis followed by Gibson Assembly. Next, we created 6 cloning vectors for the MCP-ADAR2-DD-NES and MCP-ADAR2-DD(E488Q)-NES, cloning the PCR fragments generated above into the LentiCRISPR v2 vector digested with BamHI and XbaI using Gibson Assembly. All PCRs in this section were carried out using Kapa HiFi HotStart PCR Mix (Kapa Biosystems), 20 ng template and appropriate primers in 20 μl reactions. All digestions in this section were carried out in 50 μl reactions for 3 hours at 37 °C using 2 μg of plasmid and 10 units of enzyme(s). All Gibson Assembly reactions in this section were carried out using 50 ng backbone and 30 ng of insert in a 10 μl volume and incubated at 50 °C for 1 hour. Digestions and PCRs were purified using the QIAquick PCR Purification Kit (Qiagen).

#### iii. Creation of plasmid library

Once we had 6 cloning vectors corresponding to the MCP-ADAR2-DD-NES ready, we digested these with Esp3I. These digestions were carried out in 50 μl reactions for 6 hours at 37 °C using 2 μg of plasmid and 10 units of enzyme followed by heat inactivation at 65 °C for 20 minutes. The digestion reaction was then purified using the QIAquick PCR Purification Kit (Qiagen). This was followed by cloning of the 6 oligonucleotide pools into their respective cloning vectors via Gibson Assembly using 50 ng of the digested backbone and 10 ng of the purified oligonucleotide PCR products in a 10 μl reaction, incubated at 50 °C for 80 minutes. The Gibson Assembly reaction was purified by dialysis and used to electroporate ElectroMAX Stbl4 cells (ThermoFisher) as per the manufacturer’s instructions. A small fraction (1–10 μl) of cultures was spread on carbenicillin LB plates to calculate the library coverage, and the rest of the cultures were amplified overnight in 150 ml LB medium containing carbenicillin. A library coverage of at least 400x was ensured before proceeding. Plasmid libraries were sequenced using the MiSeq (300 bp PE run).

#### iv. Creation of MS2-adRNA vectors

We began by replacing the Cas9-P2A-Puromycin from the LentiCRISPR v2 with a mCherry-P2A-Hygromycin by digesting the backbone with XbaI and PmeI. We used fusion PCRs to create the mCherry-P2A-Hygromycin-WPRE-3’LTR(Delta U3) insert which was then cloned into the digested backbone via Gibson Assembly. We used PCRs to create a MS2-adRNA-mU6-MS2-adRNA cassette which was cloned into the Esp3I digested backbone via Gibson Assembly. 4 vectors with 2x MS2-adRNAs were created targeting 5’ and 3’ TAG and GAC. All PCRs in this section were carried out using Kapa HiFi HotStart PCR Mix (Kapa Biosystems) in 20 μl reactions. All digestions in the section were carried out in 50 μl reactions for 3 hours at 37 °C using 2 μg of plasmid and 10 units of enzymes. All Gibson Assembly reactions in this section were carried out using 50 ng backbone and 20-40 ng of insert in a 10 μl volume and incubated at 50 °C for 1 hour. Digestions and PCRs were purified using the QIAquick PCR Purification Kit (Qiagen).

#### v. Lentivirus production

HEK293FT cells were maintained in DMEM supplemented with 10% FBS (Thermo Fisher) and 1% Antibiotic-Antimycotic (Thermo Fisher) in an incubator at 37 °C and 5% CO_2_ atmosphere. To produce lentivirus particles, HEK293FT cells were seeded in 15-cm tissue culture dishes 1 day before transfection and were 60% confluent at the time of transfection. Before transfection, the culture medium was changed to prewarmed DMEM supplemented with 10% FBS. For each 15-cm dish, 36 μl of Lipofectamine 2000 (Thermo Fisher) was diluted in 1.2 ml OptiMEM (Thermo Fisher). Separately, 3 μg pMD2.G (gift from Didier Trono, Addgene #12259), 12 μg of pCMV delta R8.2 (gift from Didier Trono, Addgene #12263) and 9 μg of lentiviral vector were diluted in 1.2 ml OptiMEM. After incubation for 5 min, the Lipofectamine 2000 mixture and DNA mixture were combined and incubated at room temperature for 30 minutes. The mixture was then added dropwise to HEK293FT cells. Viral particles were harvested 48 h and 72 h after transfection, further concentrated to a final volume of 500-1000 μl using 100 kDA filters (Millipore), divided into aliquots and frozen at −80 °C. Lentivirus was produced individually for all MS2-adRNA vectors and in a pooled format for the libraries. While producing lentivirus, libraries were grouped together as 1+2,3,4,5+6 so as to facilitate sequencing using the NovaSeq 6000 (250 bp PE run).

#### vi. Creation of a clonal cell line with MS2-adRNA

HEK293FT cells grown in a 6-well plate were transduced with lentiviruses (high MOI) carrying 2x MS2-adRNA targeting 5’ and 3’ TAG and GAC to create 4 different cell lines. For transductions, the lentivirus was mixed with DMEM supplemented with 10% FBS (Thermo Fisher) and Polybrene Transfection reagent (Millipore) at a concentration of 5 μg/ml and added to HEK293FT cells at 40-50% confluency. Hygromycin (Thermo Fisher) was added to the media at a concentration of 100 μg/ml, 48 hours post transduction. Top 1% of mCherry expressing cells for each line were then sorted into a 96 well plate. 3 clones of each of the 4 cell lines were then frozen down.

#### vii. Screen

Lentiviral libraries 1+2 and 3 were used to transduce clones with the 5’ TAG and GAC MS2-adRNA and libraries 4 and 5+6 were used to transduce clones with the 3’ TAG and GAC MS2-adRNA stably integrated. Transductions were carried out in duplicates. The lentiviral libraries were mixed with DMEM supplemented with 10% FBS (Thermo Fisher), Hygromycin (Thermo Fisher) at 100 μg/ml, Polybrene Transfection reagent (Millipore) at a concentration of 5 μg/ml and added to the stable clones harboring the MS2-adRNA in a 15 cm dish at 40-50% confluency. To ensure most cells received 0 or 1 ADAR2 variant, cells were transduced at a low MOI of 0.2-0.4. 24 hours post transfections, cells were passaged 1:4 into a new 15 cm dish and grown in DMEM supplemented with 10% FBS (Thermo Fisher) and Hygromycin (Thermo Fisher) at 100 μg/ml. 48 hours post transductions, the growth medium was changed to DMEM supplemented with 10% FBS (Thermo Fisher) and Puromycin (Thermo Fisher) at 3 μg/ml. 72 hours post transduction, fresh growth medium with Puromycin was added to the cells. 96 hours post transductions, the growth media was taken off and cells were washed with PBS and then harvested. Cell pellets were stored at −80 °C until RNA extraction. At least 1000x coverage was maintained at all steps of the screen.

#### viii. RNA, cDNA, amplifications, indexing

RNA was extracted using the RNeasy mini kit (Qiagen) as per the manufacturer’s instructions. cDNA was synthesized from RNA using the Protoscript II First Strand cDNA synthesis Kit (NEB). To ensure library coverage of 500x, 5 ng of RNA was converted to cDNA per library element in every sample of the screen. The volume of each cDNA reaction was 90 μl with 4.5 μg RNA, 45 μl of the Reaction mix, 9 μl Random primers and 9 μl Enzyme. Samples were incubated in a thermocycler at 25 °C for 5 min; 42 °C for 80 min; 80 °C for 5 min. The entire volume of the cDNA reaction was used to set up PCR reactions. The volume of each PCR reaction was 100 μl with 44 μl cDNA, 6 μl primers (10 μM) and 50 μl Q5 high fidelity master mix (NEB). The thermocycling parameters were: 98 °C for 30 s; 24-28 cycles of 98 °C for 10 s, 62 °C for 15 s, and 72 °C for 35 s; and 72 °C for 2 min. The numbers of cycles were tested to ensure that they fell within the linear phase of amplification. The amplicons were 440-570 bp in length and purified using the QIAquick PCR Purification Kit (Qiagen). To continue maintaining at least 500x coverage, at minimum 0.15 ng of the PCR product per library element was used to set up a second PCR adding indices onto the libraries. This was done in 50 μl reactions using 3 μl dual index primers (NEB), 135 ng purified PCR product from the previous reaction and 25 μl Q5 high fidelity master mix (NEB). The thermocycling parameters were: 98 °C for 30 s; 5-8 cycles of 98 °C for 10 s, 65 °C for 20 s, 72 °C for 35 s; and 72 °C for 2 min. The numbers of cycles were tested to ensure that they fell within the linear phase of amplification. Amplicons were purified with Agencourt AMPure XP beads (Beckman Coulter) at a 0.8 ratio. The libraries were quantified using the Qubit dsDNA HS assay kit (Thermo Fisher) and pooled together at a concentration of 10 nM for sequencing on a 250 bp PE run on the NovaSeq 6000.

#### ix. Sequencing analysis

Raw fastq reads were aligned to the ADAR2 reference sequence using minimap2 (*48*) in short-read mode with default parameters. For libraries with overlapping paired end reads, the reads were first combined using FLASH (*49*). The aligned reads were then classified into library members using strict filtering, i.e. reads were only included if they perfectly matched exactly one library member, aside from the target ADAR editing site. The editing rate at this target site was then quantified for each library member and averaged across two replicates with weights for differential coverage. To analyze the degree to which each library member differed in editing rate from the wild-type, we performed a two-proportion Z-test using a pooled sample proportion to calculate the standard error of the sampling distribution, and a two-tailed procedure to calculate p-values. Note that the wild-type rate was restricted to the rate measured within each library, such that each library member was compared only to the wild-type rate measured in the same biological context. Z-scores were calculated as follows, where *x* is the RNA editing rate, and *n* is the number of counts:

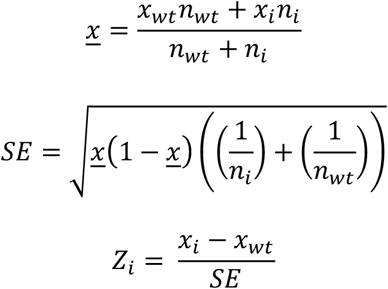

The library classification and editing quantification procedures were carried out using a custom python package. Heatmap plotting was done with modified code from Enrich2 (https://github.com/FowlerLab/Enrich2) (*50*).

#### x. Cloning individual mutants

We began by creating a cloning vector with the MCP inserted into the LentiCRISPR v2 vector digested with BamHI and XbaI using Gibson Assembly. This vector was then digested with BamHI to clone the DD mutants. All mutants were created using mutagenesis PCR followed by Gibson Assembly. All PCRs in this section were carried out using Q5 PCR Mix (NEB), 5 ng template and appropriate primers in 20 μl reactions. All digestions in this section were carried out in 50 μl reactions for 3 hours at 37 °C using 3 μg of plasmid and 20 units of enzyme(s). All Gibson Assembly reactions in this section were carried out using 30 ng backbone and 15 ng of insert in a 6 μl volume and incubated at 50 °C for 1 hour. Digestions and PCRs were purified using the QIAquick PCR Purification Kit (Qiagen).

#### xi. Luciferase assay

All HEK 293FT cells were grown in DMEM supplemented with 10% FBS and 1% Antibiotic-Antimycotic (Thermo Fisher) in an incubator at 37 °C and 5% CO_2_ atmosphere. All *in vitro* luciferase experiments for DMS validations were carried out in HEK 293FT cells seeded in 96 well plates, at 25-30% confluency, using 250 ng total plasmid and 0.5 μl of commercial transfection reagent Lipofectamine 2000 (Thermo Fisher). Specifically, every well received 100 ng of the Cluc-W85X(TAG) or Cluc-W85X(TGA) reporters, 50ng of MCP-ADAR2-DD mutants and 100ng of the MS2-adRNA plasmids. In cases where less than 3 plasmids were needed, a balancing plasmid was added to keep the total amount per well as 250 ng. 48 hours post transfections, 20 μl of supernatant from cells was added to a Costar black 96 well plate (Corning). For the readout, 50 μl of Cypridina Assay buffer was mixed with 0.5 μl Vargulin substrate (Thermo Fisher) respectively and added to the 96 well plate in the dark. The luminescence was read within 10 minutes on Spectramax i3x or iD3 plate readers (Molecular Devices) with the following settings: 5 s mix before read, 5 s integration time, 1 mm read height.

#### xii. RNA editing

RNA editing experiments for targeting *5’-GA-3’* were carried out in HEK 293FT cells seeded in 24 well plates using 1000ng total plasmid and 2ul of commercial transfection reagent Lipofectamine 2000 (Thermo Fisher). Specifically, every well received 500 ng each MCP-ADAR2-DD fragments and the adRNA plasmids. Cells were transfected at 25-30% confluence and harvested 48 hours post transfection for quantification of editing. RNA from cells was extracted using the RNeasy Mini Kit (Qiagen). cDNA was synthesized from 500ng RNA using the Protoscript II First Strand cDNA synthesis Kit (NEB). 1ul of cDNA was amplified by PCR with primers that amplify about 200 bp surrounding the sites of interest using OneTaq PCR Mix (NEB). The numbers of cycles were tested to ensure that they fell within the linear phase of amplification. PCR products were purified using a PCR Purification Kit (Qiagen) and sent out for Sanger sequencing. The RNA editing efficiency was quantified using the ratio of peak heights G/(A+G).

### Split-ADAR2

#### i. Vector design and construction

We began by digesting the pAAV_hU6_mU6_CMV_GFP with AflII to clone the NES-FLAG-MCP-linker and linker-4xλN-HA-NES downstream of the CMV promoter which were amplified from the MCP-ADAR2-DD-NLS (*23*) and 4x-λN-cdADAR2 (*14*) respectively. AvrII digestion sites were included downstream of the NES-FLAG-MCP-linker and upstream of the linker-4xλN-HA-NES to facilitate cloning of the split fragments. All split fragments were amplified from the MCP-ADAR2-DD-NLS or MCP-ADAR2-DD(E488Q)-NLS (*23*). For each split-ADAR2 pair, the N-terminal DD fragment was cloned downstream of the NES-FLAG-MCP-linker and the C-terminal DD fragment was cloned upstream of the linker-4xλN-HA-NES using Gibson Assembly. MS2-MS2, MS2-BoxB, BoxB-MS2 and BoxB-BoxB adRNA were created by annealing primers and cloned downstream of the hU6 promoter into the AgeI+NheI digested pAAV_hU6_mU6_CMV_GFP using Gibson Assembly. All PCRs in this section were carried out using Kapa HiFi HotStart PCR Mix (Kapa Biosystems) in 20 μl reactions. All digestions in this section were carried out in 50 μl reactions for 3 hours at 37 °C using 3 μg of plasmid and 20 units of enzyme(s). All Gibson Assembly reactions in this section were carried out using 40 ng backbone and 5-20 ng of insert in a 10 μl volume and incubated at 50 °C for 1 hour. Digestions and PCRs were purified using the QIAquick PCR Purification Kit (Qiagen).

#### ii. Luciferase assay

All HEK 293FT cells were grown in DMEM supplemented with 10% FBS and 1% Antibiotic-Antimycotic (Thermo Fisher) in an incubator at 37 °C and 5% CO_2_ atmosphere. All *in vitro* luciferase experiments for the split-ADAR2 were carried out in HEK 293FT cells seeded in 96 well plates, at 25-30% confluency, using 400 ng total plasmid and 0.6 μl of commercial transfection reagent Lipofectamine 2000 (Thermo Fisher). Specifically, every well received 100 ng each of the Cluc-W85X(TAG) reporter, N- and C-terminal ADAR2 fragments and the adRNA plasmids. In cases where less than 4 plasmids were needed, a balancing plasmid was added to keep the total amount per well as 400 ng. 48 hours post transfections, 20 μl of supernatant from cells was added to a Costar black 96 well plate (Corning). For the readout, 50 μl of Cypridina Glow Assay buffer was mixed with 0.5 μl Vargulin substrate (Thermo Fisher) and added to the 96 well plate in the dark. The luminescence was read within 10 minutes on Spectramax i3x or iD3 plate readers (Molecular Devices) with the following settings: 5 s mix before read, 5 s integration time, 1 mm read height.

#### iii. RNA editing

All *in vitro* RNA editing experiments were carried out in HEK 293FT cells seeded in 24 well plates using 1500ng total plasmid and 2ul of commercial transfection reagent Lipofectamine 2000 (Thermo Fisher). Specifically, every well received 500 ng each of the N- and C-terminal ADAR2 fragments and the adRNA plasmids. In cases where less than 3 plasmids were needed, a balancing plasmid was added to keep the total amount per well as 1500 ng. Cells were transfected at 25-30% confluence and harvested 48 hours post transfection for quantification of editing. RNA from cells was extracted using the RNeasy Mini Kit (Qiagen). cDNA was synthesized from 500ng RNA using the Protoscript II First Strand cDNA synthesis Kit (NEB). 1ul of cDNA was amplified by PCR with primers that amplify about 200 bp surrounding the sites of interest using OneTaq PCR Mix (NEB). The numbers of cycles were tested to ensure that they fell within the linear phase of amplification. PCR products were purified using a PCR Purification Kit (Qiagen) and sent out for Sanger sequencing. The RNA editing efficiency was quantified using the ratio of peak heights G/(A+G). RNA-seq libraries were prepared from 250ng of RNA, using the NEBNext Poly(A) mRNA magnetic isolation module and NEBNext Ultra RNA Library Prep Kit for Illumina. Samples were pooled and loaded on an Illumina Novaseq (100 bp paired-end run) to obtain 40-45 million reads per sample.

#### iv. Quantification of RNA-seq A-to-G editing

RNA-seq analysis for quantification of transcriptome-wide A-to-G editing was carried out as described in (*23*).

